# CD4+ T cell subsets present stable relationships in their T cell receptor repertoires

**DOI:** 10.1101/2020.11.01.364224

**Authors:** Shiyu Wang, Longlong Wang, Ya Liu

## Abstract

CD4+ T cells are key components of adaptive immunity. The cell differentiation equips CD4+ T cells with new functions. However, the effect of cell differentiation on T cell receptor (TCR) repertoire is not investigated. Here, we examined the features of TCR beta (TCRB) repertoire of the top clones within naïve, memory and regular T cell (Treg) subsets: repertoire structure, gene usage, length distribution and sequence composition. First, we found that memory subsets and Treg would be discriminated from naïve by the features of TCRB repertoire. Second, we found that the correlations between the features of memory subsets and naïve were positively related to differentiation levels of memory subsets. Third, we found that public clones presented a reduced proportion and a skewed sequence composition in differentiated subsets. Furthermore, we found that public clones led naïve to recognize a broader spectrum of antigens than other subsets. Our findings suggest that TCRB repertoire of CD4+ T cell subsets is skewed in a differentiation-depended manner. Our findings show that the variations of public clones contribute to these changes. Our findings indicate that the reduce of public clones in differentiation trim the antigen specificity of CD4+ T cells. The study unveils the physiological effect of memory formation and facilitates the selection of proper CD4+ subset for cellular therapy.

## Introduction

CD4+ T cells play critical roles in mediating adaptive immunity. Via T cell receptors (TCR), CD4+ T cells recognize the complex of epitopes and major histocompatibility complex II and then induce the activation of other cells in infections^1,2^, cancer^3^ and autoimmune diseases.

To acquire mature functions, CD4+ T cells undergo differentiation. NT is the protype of CD4+ T cell and has the greatest potential among CD4+ T subsets to differentiate to other subsets. NT usually keep a serenity and can refresh themselves by proliferation. When NT encounters pathogens, it will home to lymphatic organs and receive the help from dendritic cells to initiate the polarization. The study on TCR repertoire suggests that NT has the most large scale of evenness of TCR repertoire among all CD4+ subsets^4^, which indicates the greatest potential to recognize antigens. In a classical differentiation model^5,6^, naive (NT) senses stimulations via TCR, polarizes and then differentiates to effector T (ET). ET plays the key role to mediate adaptive immune response, although the amount of ET in peripheral blood is limited. After few weeks’ activation, a small part of ET differentiates to memory. Memory is sensitive to antigens, while always keeps silent. In peripheral blood, central memory (CMT), effector memory (EMT) and stem-cell like memory T cell (Tscm) are main subsets of memory. CMT and Tscm have a potential to be self-renewal and is found to affect the infections. EMT, compared to CMT^7^, is long-lived and has a lower threshold to reactivate to pathogens. Compared to NT, memory subsets have a lower diversity of TCRB repertoire^8^. However, it is unclear that large differences between NT and memory subsets exist in the functions of TCR repertoire or not. Treg is a special subset of CD4+ cell and usually plays a role to regulate functions, proliferation, and differentiation of conventional T cells^9^.

TCR plays the key role to determine T cell functions^10^. For NT, signals via diverse TCRs reform the TCR repertoire, and skew their differentiation potential. For instance, strong TCR signals during viral infection correspond to helper T cell differentiation, and comparative lower signals facilitate the differentiation of memory and follicular T cell^11^. Memory’s TCR repertoire composition determines the possibility to provide a rapid protection for individuals to against former and, sometimes, novel antigens. For example, the architecture of the TCR repertoire contributes to the performance of the adaptive immune response against pathogens, such as SARS-CoV-2^12^. Cross-reactivation from memory can provide a rapid protection to a novel pathogen in some individuals, such as the case reports of COVID-19^13^. The importance of TCR repertoire for memory cell functions was found in tissues, where the differential composition of TCR repertoire of CD4+ memory among tissues equipped them with distinct functions^14^. The function of Treg was restricted by TCR repertoire. The optimal diversity of TCR was essential for the suppressive ability^15^, and limitations on TCR diversity disturbed the self-tolerance of immune system^16^. Although evidences show that the features of TCR repertoire are distinct among CD4+ T subsets, the effect of differentiation on T cell receptor (TCR) repertoire of CD4+ T cells are not investigated.

To unveil the influence of differentiation on TCR repertoire, we analyzed the sequencing data of TCR beta (TCRB) chain of NT, ET, EMT, CMT, Tscm and Treg. We detected repertoire structure, germline gene usage, sequence composition and public clones of TCRB repertoire of each subset. We found that NT, CMT, Tscm and Treg were discriminated from each other by repertoire structure, gene usage and sequence composition, independently. The TCRB repertoires of NT are similar to the TCRB repertoires of less-differentiated memory subset (for example, Tscm and CMT). The TCRB repertoires of ET and the TCRB repertoires of EMT are sensitive to the heathy state and have flexible relationships with other subsets. Public clones account for the main part of top clones of NT and reduce along the CD4+ cell differentiation. The enrichment of public clones shortens the length distribution of top clones in NT. Public clones are polyfunctional and broaden the antigen spectrum recognized by NT. Our findings disclose the differential functions of CD4+ cell subsets and the influence of differentiation on TCR repertoire. Our findings facilitate the selection of CD4+ subsets for cellular therapy.

## Material and Methods

### Datasets

In this study, we conducted analyses on high-throughput TCR repertoire datasets for CD4+ T cell subsets from publication described as follows.

Dataset1 included Naïve (NT), central memory (CMT), stem-cell like memory (Tscm) and Treg from eight healthy individuals and eight type-one-diabetes (T1D) patients ^17^. CD4+ T cells were sorted into subsets by fluorescence-activated cell sorting (FACS), and then RNA was extracted and sequenced in parallel. For validation, we employed dataset2 of five CD4+ T cell subsets: NT, effector (ET), CMT, effector memory (EMT) and Treg from other ten rheumatoid arthritis patients (RA)^18^. Top1000 clones are referred as the 1000 clones with the highest frequency. The V-/J-segments used by top1000 clones were extracted for statistic.

### Statistical analysis and plots

Statistical analyses were performed with R. Significance was examined by Willcox ranked test. The correlation coefficients and significance was tested using cor.test() in R with default parameters. Graphics were generated with R package ggplot2. Principle component analysis (PCA) was conducted with R package *forcats*. Data was treated with R package *readr, dplyr* and *tidyr*.

### Definition of a clone

For all analyses, clones were defined as the amino acid sequence identity of CDR3 (TCRB) regions. CDR3s from dataset1 ware defined and annotated by IMonitor^19^, and CDR3s from dataset2 were reported by authors.

### Determination of diversity

Renyi entropy was used in our study to evaluate the diversity with alpha value from 0 to 20. When alpha is 1, the index equals to the Shannon index. Renyi entropy formula is 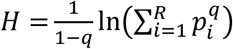, and Shannon diversity index formula is 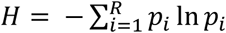, where *H* is the diversity index, *q* is alpha value, *R* is the total number of clones for analysis, and *p*_*i*_ is the frequency of the *i*th clone.

### Determination of Jessen-Shannon distance

Jessen-Shannon distance (JSD) is used to evaluate the similarity in repertoire architecture among subsets^20^. We calculated JSD with JSD(), a function included in philentropy ()^21^, a R package. A low JSD indicates that TCRB repertoire structures are similar.

### Identify the contribution of k-mer to PCA classification

We used prcomp() to calculate the principle components (PC) for data, and estimated the contribution of each k-mer to PC1 and PC2 with cos2().

### KeBABS SVM analysis

SVM analysis was performed using kernel-based analysis of biological sequences with a R package KeBABS^22^. Amino acid sequence of clones was split into features with length k = 3, and cost parameter C = 100 was used for the misclassification of a sequence. For all SVM analyses, data was split into training (80%) and test (20%) set. SVM training was performed on the training set, and class prediction was performed on the test set. Prediction accuracy of classification was qualified by calculating 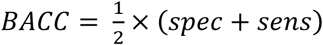, where specificity was calculated as 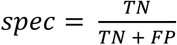, and sensitivity was defined as 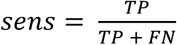, (where TN = true negative, FP = false negative, TP = true positive and FN = false negative). The area under the receiver operating characteristic curve (AUC) was calculated, where the AUC = 0.5 means a random classification (BACC = 50%), and AUC = 1 means a perfect classification (BACC = 100%).

### Determination of epitope specificity of clones by GLIPH2

GLIPH2^23^ is a robust tool to predict the cluster of clones targeting the same epitope. Here, we use this method to unveil the diversity of potential epitopes targeted by top1000 clones in each subset. The reference of CD4+ T clones and their gene usage, the distribution of gene usage of CD4+ T and length distribution of CDR3 were included in ref_CD4.txt, ref_V_CD4.txt and ref_L_CD4.txt. These reference files were downloaded from the official website of GLIPH2 (http://50.255.35.37:8080/). A filter with a high stringency (Fisher_score < 0.0001, number_subject >= 3 and number_unique_cdr3 >= 3) was used to improve the prediction accuracy.

## Results

### The TCRB repertoire structure of NT is similar to The TCRB repertoire structure of CMT and Tscm

Frequent clones affect the immune repertoire structure^24^. We thus performed the analyses on top1000 clones within each subset. Renyi entropy with alpha values from zero to twenty was used to evaluate the diversity. In dataset1, The TCRB repertoire of NT and Tscm present similar diversities at all alpha values, and are more diverse than the TCRB repertoire of CMT and the TCRB repertoire of Treg (Figure 1A). In dataset2, NT has the most diverse TCRB repertoire among all subset whereas ET has the lowest. The TCRB repertoire of CMT is more diverse than that of ETM and Treg. (Supplementary figure 1A). The similarity of TCRB repertoire structure of subsets was estimated by Jensen-Shannon distance. In dataset1, the TCRB repertoire structure of NT is similar to the TCRB repertoire structure of less-differentiated subsets (CMT and Tscm), but the TCR repertoire structures of Tscm and CMT are different from each other; the TCRB repertoire of Treg is different to the TCRB repertoire of NT and CMT with high JSDs. It indicates that Treg has a structure of TCRB repertoire like that of more-differentiated memory subsets (Figure 1B). In dataset2, NT and CMT have similar TCRB repertoire structures, and the TCRB repertoire structure of Treg is similar to the TCRB repertoire structure of EMT rather than that of CMT (Supplemental Figure 1B). These findings fit with the trend found in dataset1. To consider the overlapping usage of CDR3 clones, we further evaluated the similarity of TCRB repertoire among subsets with the Morisita-Horn similarity index. In this analysis, NT keeps a similar TCRB repertoire like Tscm and CMT, while the TCRB repertoire of NT is different from the TCRB repertoire of EMT and Treg (Figure 1C; Supplemental Figure 1C). In conclusion, the TCRB repertoire structure of CD4+ T cells is skewed along with cell differentiation levels.

**Figure 1.**
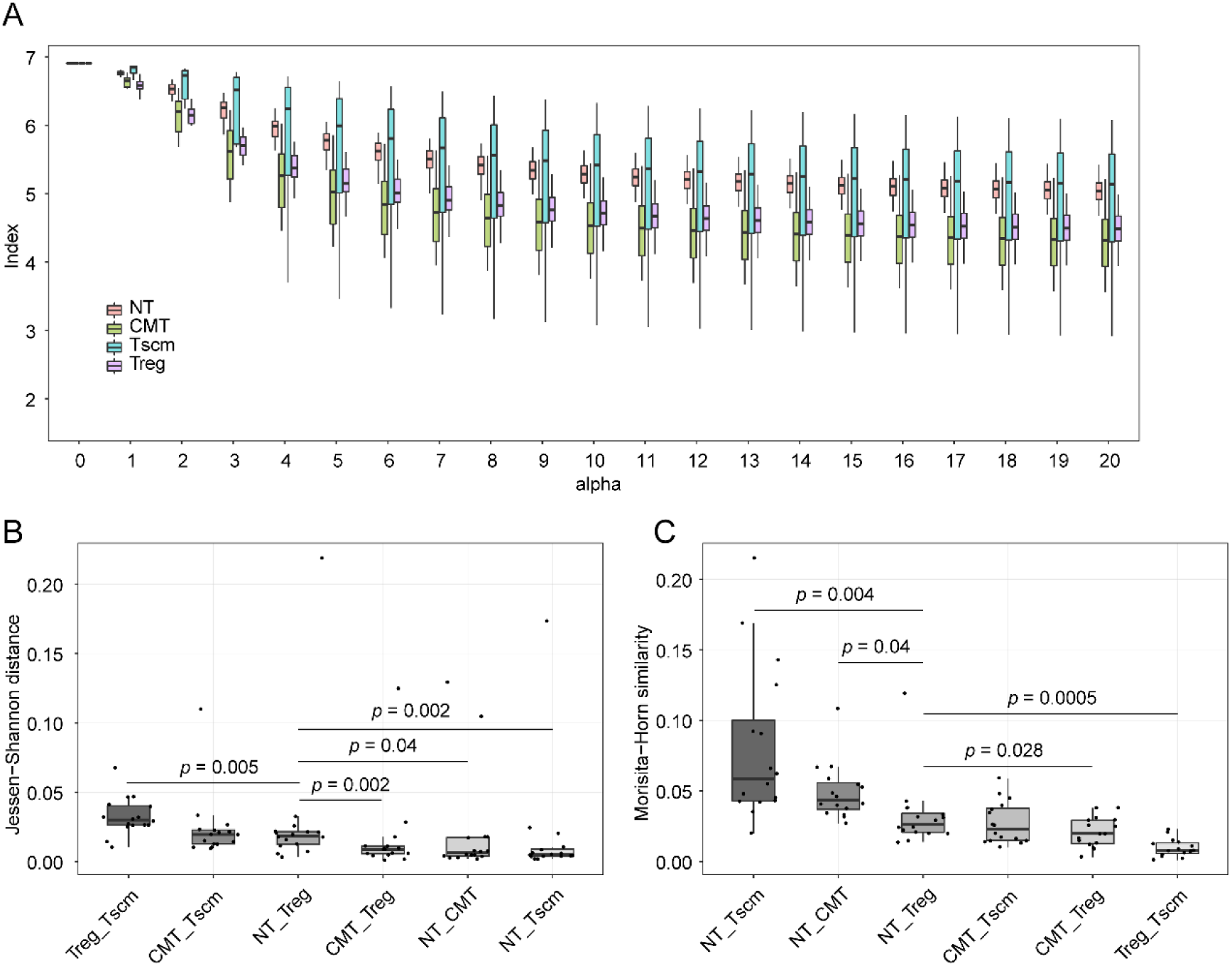
The diversity of TCRB repertoire of four subsets and the relationship of their TCRB repertoire structures. (**A**) The Renyi entropy index of all subsets with alpha value from 0 to 20. When alpha is equal to 1, the index was calculated as Shannon-index. (**B**) The Jessen-Shannon distance between subsets. The difference between NT and other subsets, and Treg and other subsets were tested. (**C**) The Morisita-Horn similarity between subsets. Paired Wilcox-ranked test was used in **B** and **C**.

### Gene usage is distinct among subsets and a part of genes are skewed along subsets

We used principle component analysis (PCA) to examine the discrimination of gene usage among subsets. NT, CMT, Tscm and Treg could be distinguished from each other by V- and J-gene respectively (Figure 2A and B; Supplemental Figure 2A). However, EMT and ET could not be separated from other subsets in dataset2 (Supplemental Figure 2A). We examined the genes usage of each subset, and found that 24 of 72 genes were differently used by NT, Tscm and CMT in dataset1, 23 of 72 genes were differently used by NT and CMT in dataset2. There are 11 increased genes and 13 decreased genes in CMT, compared with NT in dataset1; and 3 increased genes and 20 decreased genes in dataset2, compared with NT in dataset2. Notably, these genes continuously changed according to NT, CMT, ET, EMT and Treg (Supplemental Figure 2C and D).

**Figure 2.**
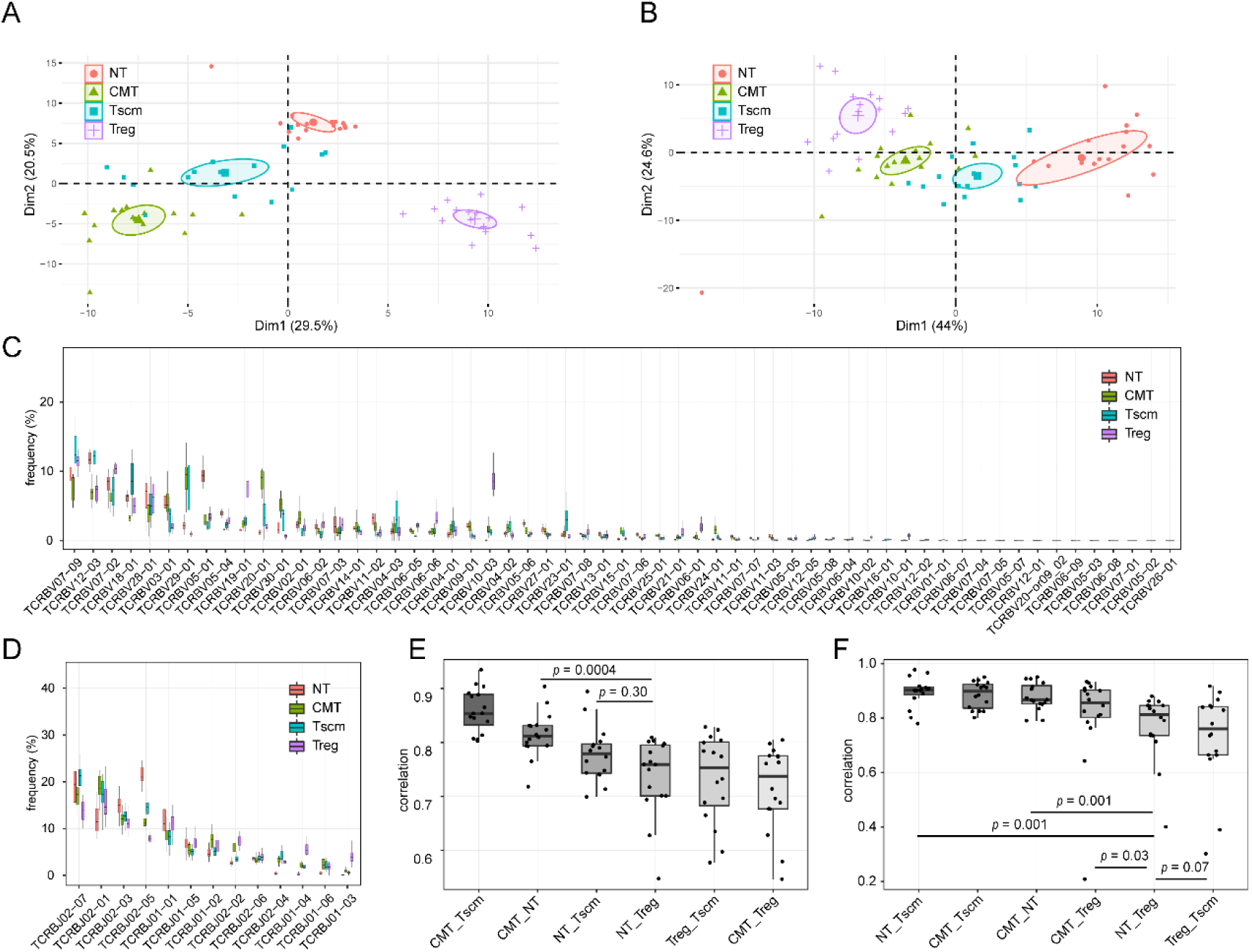
The V- and J-gene usage of CD4+ T subsets and the relationships of the gene usage of subsets. (**A**) PCA for the frequency of V-genes of each subset. (**B**) PCA for the frequency of J-genes of each subset. (**C**) The frequency of V-genes of each subset. (**D**) the frequency of J-gene within each subset. (**E**) The Spearman correlation of V-gene usage between subsets. (**F**) The Spearman correlation of J-gene usage between subsets. Paired Wilcox-ranked test was used in **E** and **F**.

With the correlations of V-gene usage among subsets, we found that NT, CMT and Tscm show high correlations with each other, while their correlations with Treg are low (Figure 2E; Supplementary Figure 2E). For J-gene, NT also shows a high correlation with CMT and Tscm, whereas a low correlation with Treg. In dataset2, the V-gene usage of NT highly correlates to the V-gene usage of CMT rather than EMT and Treg (Supplementary Figure 2F). Notably, the V-gene usage of Treg is similar to the V-gene usage of NT rather than to the V-gene usage of CMT in the T1D donors (p=0.0005, Supplementary Figure 4) but not in health donors (p=0.05). This phenomenon suggests Treg has a flexible relationship with NT depending on the heathy states of donors. In conclusion, NT, memory subsets and Treg have distinct gene usages, and the gene usage of NT is more similar to less-differentiated memory subsets (CMT and Tscm) rather than EMT.

### CDR3 sequence composition are different among TCRB repertoire of subsets and indels contribute to these differences highly

We examined the CDR3 sequence composition by decomposing kernels containing three amino acids. To identify the difference in the composition among subsets, we used PCA to discriminate subsets. In PCA plot, subsets are distinguished from each other. Treg is close to EMT, CMT mix with ET. NT has a clear boundary to others (Figure 3A; Supplemental Figure 5A). To evaluate the correlation of k-mer among subsets, we used Spearman correlation method. The k-mer usage of NT exhibits weakened correlations with others according to Tscm, CMT and Treg in dataset1, and CMT, ET, EMT and Treg in dataset2 (Figure 3B; Supplemental Figure 5B).

**Figure 3.**
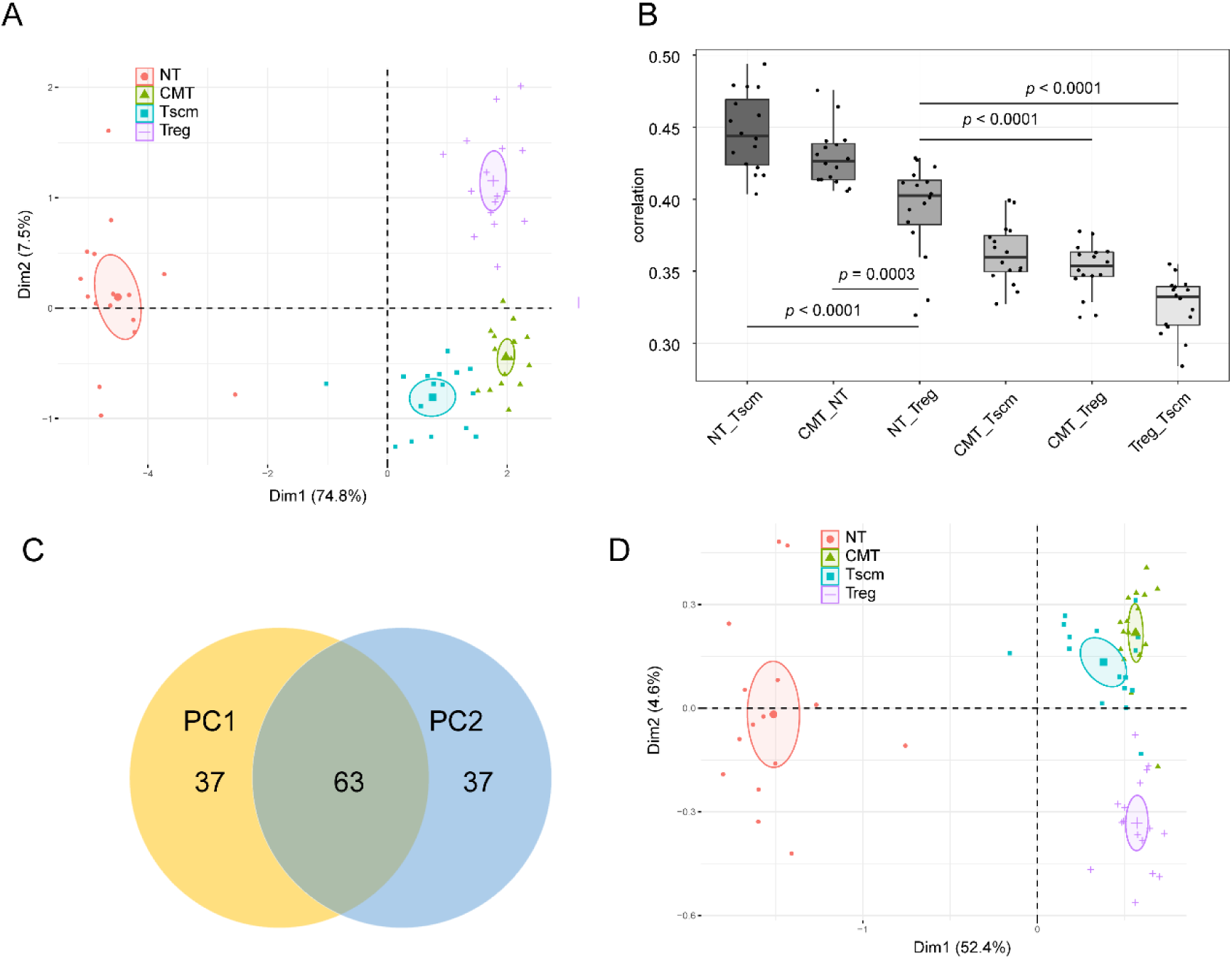
The relationship of sequence composition among subsets. (**A**) PCA based on the sequence composition for samples. (**B**) Spearman correlations of k-mer usage between subsets. (**C**) The overlap of top100 k-mers which mostly contribute to principle component 1 (PC1) and PC2. (**D**) the PCA based on sequence composition without those located on V- and J-segments for samples. Paired Wilcox-ranked test was used in **B**.

To identify the subregions where the k-mers contributes to PCA, we extracted the top100 k-mers by ranking their contributions to principle component 1 (PC1) and principle component 2 (PC2). 6371 unique k-mers were used by both of PC1 and PC2. We ranked k-mers by their contributions to PC1 and PC2 respectively, and found that PC1 and PC2 shared 63 top100 k-mers (Figure 3C; Supplemental Figure 5C). Then we aligned k-mer to references, and showed that 29 top100 k-mers located in V-region and 47 in J-region. However, after we removed the k-mer enriching in V-/J-segments, we found that NT, Tscm, CMT and Treg still kept distinctions to each other (Figure 3D). we further perform a same operation on dataset2 to remove the k-mer in V- and J-segments, and showed a similar result that NT, CMT and Treg are able to be distinguished from each other (Supplemental Figure 5D). Since the gene usage, insertion and deletion (indel) are factors to skew the CDR3 sequence compositions^25^, these results suggests that indel in N1-D-N2 region rather than gene usage contribute to the difference of sequence composition among subsets.

### Top1000 clones within NT are shorter than those in other subsets and the shortness is little affected by V-/J-gene usage

The entire repertoire of NT was reported to be longer than the repertoire of memory^26^. However, we found that the top clones in NT were shorter than clones in other subsets in all datasets (Figure 4A; Supplemental Figure 6). Via calculation of the Pearson correlations, the length distribution of NT is different to the length distribution of CMT and Tscm. It suggests that the length distribution of TCRB repertoire of NT is highly skewed in these less-differentiated memory cells (Figure 4B). Since the naïve cells are sorted without antibody against CD27 in dataset2, the length distribution of NT can be affected by the contamination from cell sorting. We examined the length distribution in dataset 1.

**Figure 4.**
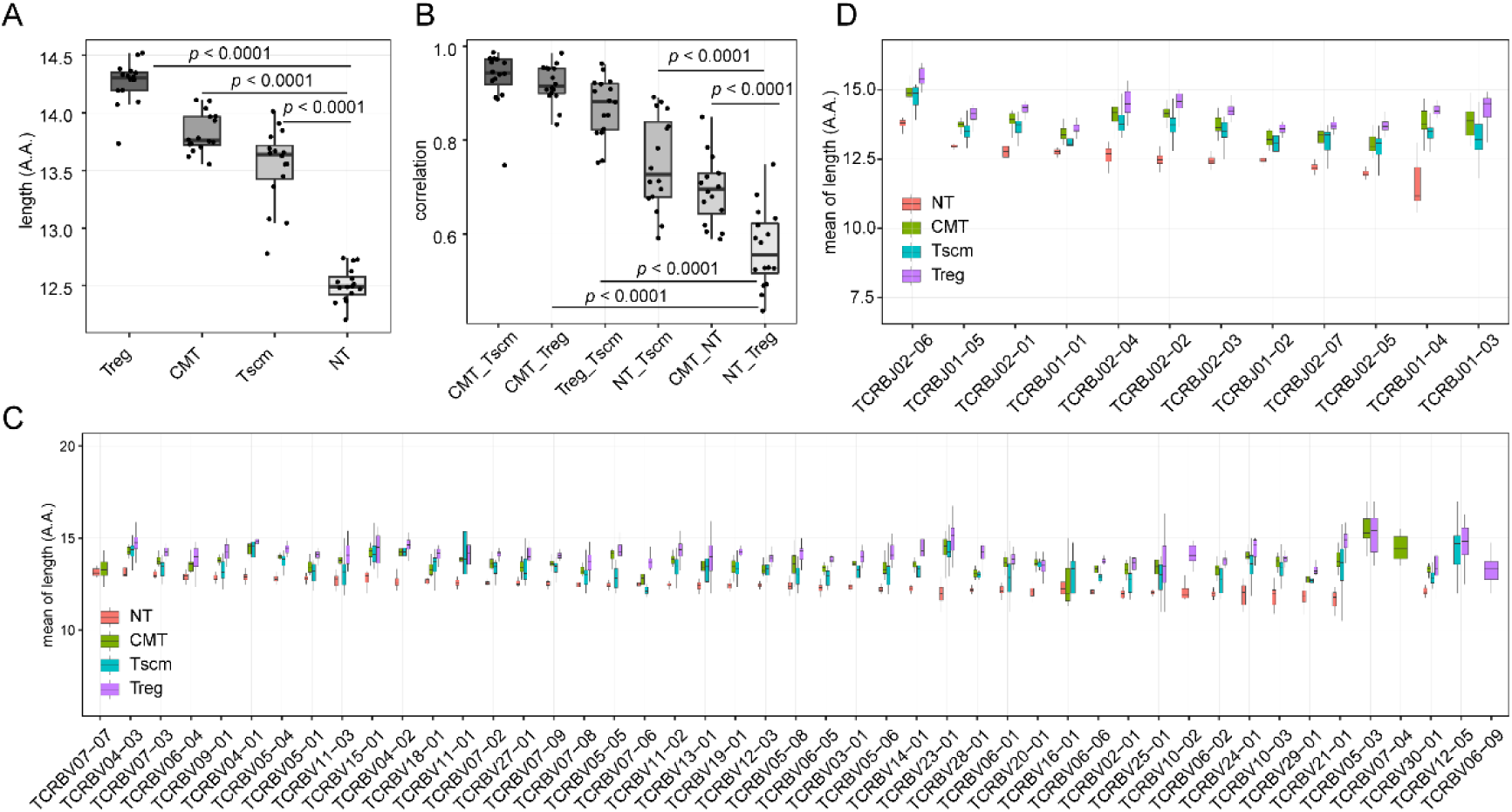
The correlations of length distribution among subsets and the influence of gene usage on length distribution. (**A**) The length distribution of CDR3s of each subset. (**B**) The correlation of length distribution between subsets. (**C**) The mean length of clones using each V-gene. (**D**) The mean length of clones using each J-gene. Paired Wilcox-ranked test was used in **A** and **B**.

To identify whether the gene usage affects the CDR3 length distribution, we calculated the mean length of clones for each gene. The mean length is different among clones by varied V- and J-genes (Figure 4C and D), however, clones using all of genes are shorter in NT than - clones in CMT and Tscm. Therefore, for top clones, the clones of NT are shorter than the clones of other subsets, and the gene usage contributes less to the distinct length distributions among subsets.

### Public clones enrich in NT and have more differences with private clones in memory subsets than in NT

Public clones that are shared by individuals were shown to be different from private clones in sequence composition^25^. Our analyses showed that public clones were shorter than private ones within top1000 clones (Supplemental Figure 7A). It suggests that public clones may affect the features of top clones. We referred clones found in no less than two individuals as public clones. We found more public clones in NT than in other subsets: 1,400 in NT, 262 in Tscm, 113 in CMT, and 72 in Treg from HD; 1544 in NT, 146 in Tscm, 128 in CMT and 92 in Treg from T1D (Figure 5A; Supplemental Figure 7B). Via calculating abundance, we found that public clones were composed of ∼70% of top1000 in NT, and ∼5% in other subsets. It suggests that the public clones in NT have a larger effect on the repertoire of top clones than the public clones in others (Figure 5B). Most of public clones in NT were a little presented in other subsets (Figure 5C), and about 50% public clones in each subset could be found in NT. This result indicates that public clones in NT are less maintained than the top clones in other subsets. To detect the differences between public clones and private clones within each subset, support vector machine (SVM) was used. To avoid the influence of differential sample sizes, we randomly down-sampled 400 public clones and private clones for each subset, respectively. The prediction was repeated for 100 times. The prediction accuracy (BACC) in NT was found to be lower than BACC in other subsets (Figure 5D). It suggests that the differences between private and public clones in differentiated subsets are larger than the differences in naïve. Further analyses showed that the gene usage of public clones is similar to the gene usage of all top1000 clones (Supplemental Figure 7C). It suggests that gene usage is not skewed in public clones.

**Figure 5.**
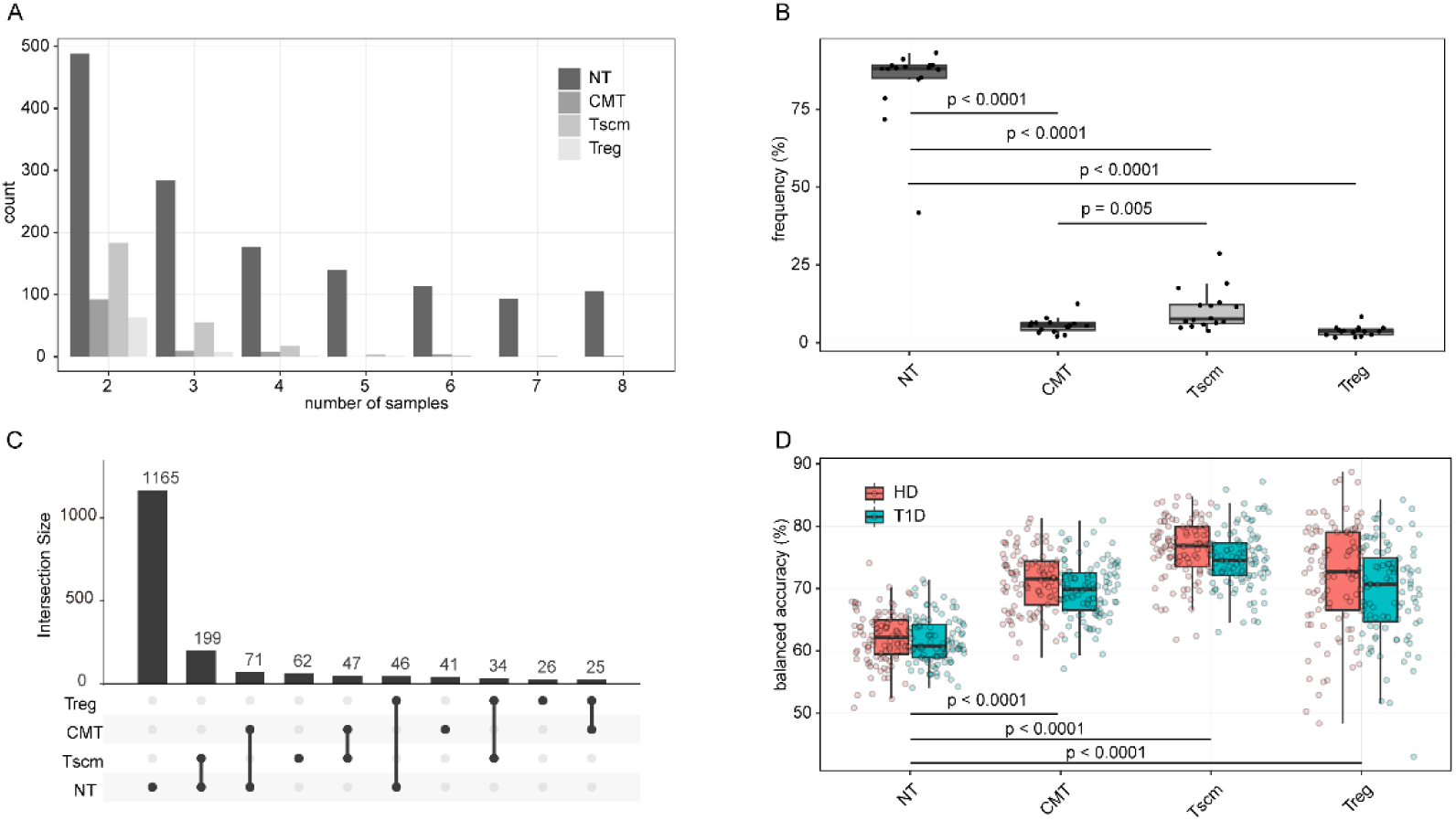
The differential usage of public clones among subsets and the discrimination of sequence composition between public clones and private clones. (**A**) The number of public clones shared by from two to eight HDs. (**B**) The percentage of unique public clones within top clones in each subset. (**C**) The overlap of public clones from HDs among subsets. (**D**) The prediction accuracy (BACC) of SVM (k = 3) for public clones and private clones based on sequence composition within each subset. Paired Wilcox-ranked test was used in **B**, Wilcox-ranked test was used in **D**.

### The sequence compositions of public clones and private clones are both skewed along differentiation

To identify that public clones or private clones account for the increased difference in memory and Treg, we performed SVM to discriminate public clones as well as private clones from different subsets separately^25^. For public clones, the BACC was from 50% to 60%; NT was able to be discriminated from Treg, CMT and Tscm with ∼ 55% BACC; whereas CMT was incapable of separating from Tscm with ∼ 50% BACC (Figure 6A). For private clones, the BACC was from 50% to 70%. We were able to achieve a high prediction accuracy to discriminate private clones of TN from private clones of Treg, but we failed to separate private clones from CMT and Tscm (Figure 6B). When we increased the sample size of private clones from 400 to 2500 for training SVM model, we found that the varied BACCs to discriminate private clones from different subsets were still existed (Supplemental Figure 8A). These results suggest that the sequence compositions of public clones and private clones are both skewed in differentiation.

**Figure 6.**
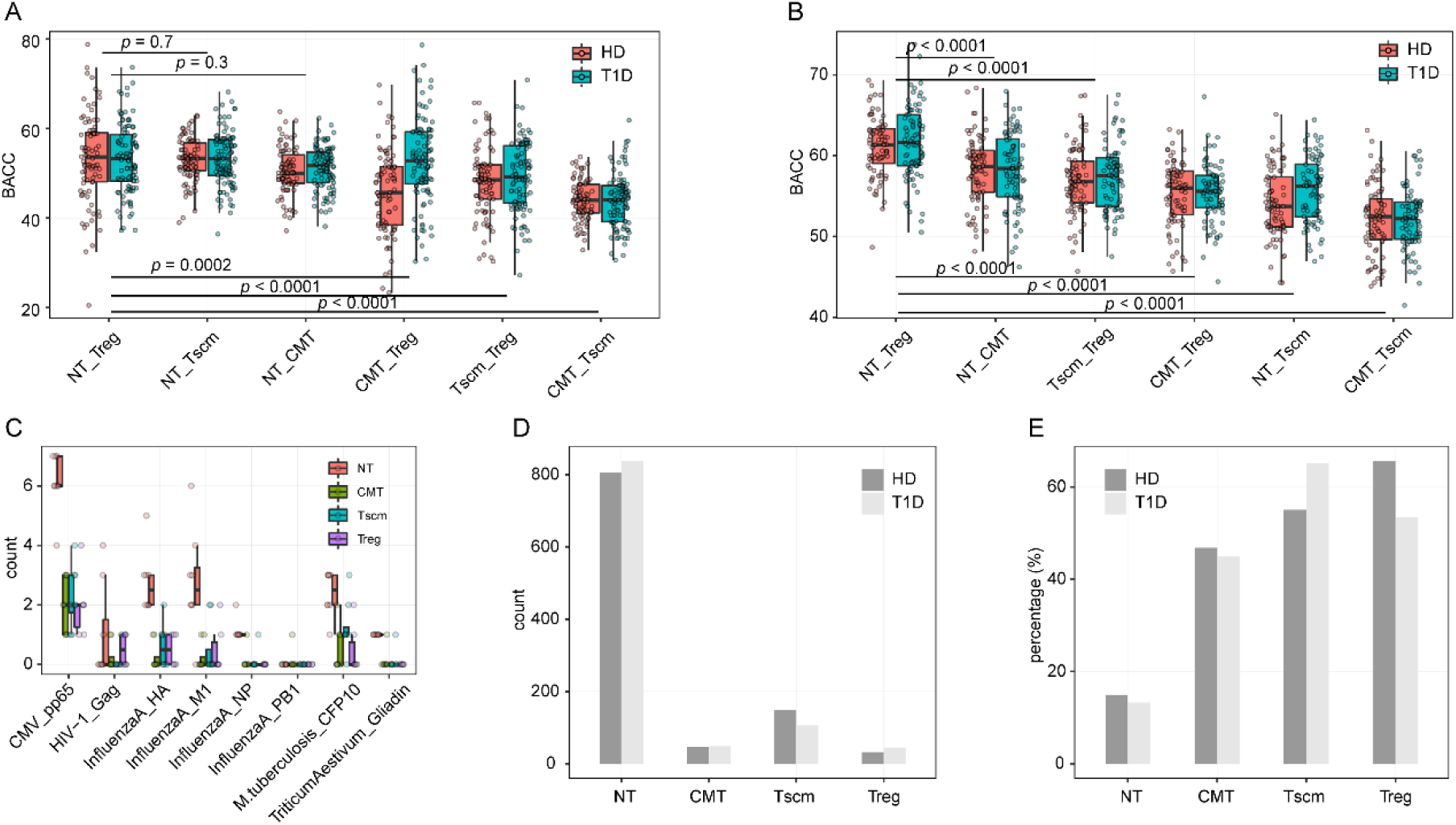
public clones maintain stable composition across subsets and provide the large part of the ability to recognize antigens. (**A**) SVM for discriminating the composition of public clones among subsets. (**B**) SVM for discriminating the composition of private clones among subsets. (**C**) The enrichment of antigen-related clones in each subset. (D) the number of clusters targeting diverse antigens predicted by GLIPH2 within each subset. (E) The percentage of clusters of private clones predicted by GLIPH2 within each subset. Wilcox-ranked test was used in **A** and **B**.

### The reduced number of public clones narrows the antigen spectrum recognized by differentiated subsets

To unveil the functions of clones among subsets, we annotated clones by VDJdb^27^. 1885 clones of CD4+ T cells targeting eight epitopes in total are recorded by this database. Comparing with CMT, Tscm and Treg, NT has more clones recognizing antigens (HA, H1 and NP) from influenza, pp65 from cytomegalovirus (CMV), CFP10 from *M. tuberculosis* and gliadin from *Triticum Aestivum* (Figure 6C; Supplementary figure 8B). To estimate the spectrum of antigens targeted by the top clones, we used GLIPH2^23^ to predict the clusters recognizing diverse antigens for each subset. With a stringency filter (see Methods), we found out 806 clusters in NT from HC and 836 clusters in NT from T1D respectively; while less than 200 clusters in whole of Tscm, CMT and Treg (Figure 6D). When public clones were removed from top clones, only 14.88% clusters remained in NT of HC and 13.25% clusters remained in T1D, whereas over 45% clusters remained in other subsets (Figure 6E). It suggests that the public clones enlarge the antigen spectrum recognized by top clones in NT. In conclusion, NT recognizes a broader antigen profile contributed by public clones.

## Discussion

It is essential for CD4+ T cells to recognize antigens with TCR, which is primarily achieved by the CDR3 region. CD4+ T cells can acquire new functions via differentiation; however, it is unclear how differentiation affects their TCR repertoire. We detected the relationships among the TCRB repertoire of top1000 clones of naïve, memory and Treg subsets (including NT, ET, Tcm, Tem, Tscm ET and Treg) by estimating the repertoire structure, the germline gene usage, the sequence composition (K-mer) and public CDR3 clone usage.

We derive that the TRBV repertoire features of memory subsets are tightly regulated in differentiation. We observed that 23 of 72 genes increased or decreased in an order of NT, Tscm, CMT and EMT. It indicates that a mechanism exists to regulate the variations across subsets. Furthermore, since Tscm is the least differentiated cell whereas EMT is the highest one among the tree memory subsets^28^, it indicates that the differentiation level is along with the mechanism. CMT is formally considered as the primary memory subset which ET prefer to differentiate to, and then part of CMT differentiates to EMT. In the past decade, Tscm has been found to mix phenotypes of naïve and memory. Tscm is able to self-renew and replenish more differentiated subsets of memory T cells, and therefore acts as the key intermediary of the generation of memory^29,30^. In together, differentiation levels of memory subsets reflect their differentiation order. However, memory cells can be directly generated from naïve cells by asymmetric cell division ^31,32^. It indicates that the differential order should not be the only factor skewing TCRB repertoire. Shown by X. L. Hou *et al*^33^, the early events in thymic T cell development are different for CD4+ naïve and memory cells. It suggests that genetic factors affect TCRB repertoire of CD4+ T cells in differentiation. In addition, events at the very early lifetime can be involved in manipulating TCRB repertoire. Observations in newborns show that memory in human develops at the very early period of lifetime^34^. Newborns less encounter pathogenic antigens. It implies that, for newborns, food, self-antigens and even cytokine driven clones compose the large part of TCRB repertoire of memory. Since the highly frequent clones in NT are self-antigen related, the features of frequent clones in NT will be delivered to memory at this period^35^. Furthermore, shown by Graeme *et al*, a half part of memory is maintained by self-renewal influx during the lifetime^36^. S. Jaafoura *et al* showed that less-differentiated memory subset is more stable during pathogen infection^28^. It suggests that Tscm and CMT rather than EMT maintain the features of TCRB repertoire inherited from NT at the early lifetime. In conclusion, it is reasonable to drive that events at the early lifetime, genetic factors and differentiation order regulate the TCRB repertoire of CD4+ T subsets with differentiated levels.

Public clones are key components that affect the features of TCRB repertoire in differentiation. First, we found that public clone usage rather than gene usage shortens the length distribution of top clones within NT. Second, the sequence composition of public clones which is skewed in differentiated subsets contribute to the variations of TCRB repertoire in differentiation. Third, decreased public clones induce a reduction in antigen spectrum recognized by memory and Treg subsets. These results suggest that the skewed public clone usage highly affect top clones in differentiated subsets. Furthermore, we showed that factors affecting the generation of public clones in memory and Treg are different to that in NT. The generation of public clones were largely attributed to genetic factors and thymic positive selection in the previous study^37^. In our study, public clones from NT are less maintained in differentiated subsets, and SVM analyses indicate that sequence composition in memory subsets is different from that of NT. These results suggest that other factors, such as antigen, trim the sequence composition of public clones. However, there are uncertainties about public clones in NT. First, it is unclear to the cause that the promiscuous ability of public clones does not induce the clonality of the public clones in effector and memory subsets. Second, it is unclear that the physiological function of public clones in NT. As a hypothesis, the promiscuous public clones in NT, sensitive to many antigens, are important to initiate the primary immune response, and this function is forbidden in differentiated subsets. Further studies with single-cell RNA-seq and paired sequencing of TCR alpha and beta chains will unveil the biological functions of the public clones in NT.

In addition to the sample size, we identified that the T cell subset also affect the SVM prediction accuracy. Shown by Victor Grief *et al*^25^, public clones and private clones can be discriminated by SVM using k-mer distribution. SVM can obtain a better prediction accuracy (BACC) by using a larger sample size. We extended the detection in NT, Tscm, CMT and Treg. By normalizing the sample size for training, we found that the prediction accuracy for public clones and private clones in Tscm, CMT and Treg is higher than the prediction accuracy in NT. It suggests that the difference between public clones and private clones is enlarged in the differentiated subsets. When we performed SVM on public clones and private clones among subsets respectively, the sequence compositions of public clones and private clones were skewed in differentiation.

A small part of peripheral Treg differentiated from conventional Treg. Shown by Golding A. *et al*, the repertoire of Foxp3+ and Foxp3-cells did not overlap^38^. Although peripheral Tregs are differentiated from conventional T cells^39^ and can introduce the features of NT into Treg, the TCR repertoire of effector and memory subsets is similar to NT than to Treg. This phenomenon suggests that the influx from naïve just composed a minor part of Treg in blood, and comparing to Treg, the features of naïve are maintained in effector and memory subsets in the differentiation. Our study includes samples of three healthy states (heathy, RA and T1D individuals), and therefore highlights that our findings are consistent in heathy conditions and datasets.

## Acknowledgments

The authors would like to thank Z.Q. Ding from Singapore institute of technology for edition of the manuscript; Y. Liu from BGI-Shenzhen and W. Zhang from department of computer science, City University of Hong Kong for comments on this manuscript.

## Conflicts of interest

The authors declare no conflicts of interest.

## Author contributions

SY. W. designed the study, performed the analyses, and drafted the manuscript; Y. L. supervised the study, and revised the manuscript.

## Financial support

This work was supported by BGI-Shenzhen, China National GeneBank (CNGB), Science, Technology and Innovation Commission of Shenzhen Municipality under grant No. JCYJ20170817145845968, and Shenzhen Key Laboratory of Single-Cell Omics (NO. ZDSYS20190902093613831).

## Notes

### Competing Interest Statement

The authors have declared no competing interest.

