## Supplemental figures for "CD4+ T cell subsets present stable relationships in their T cell receptor repertoires": SupplementrayFigures.pdf

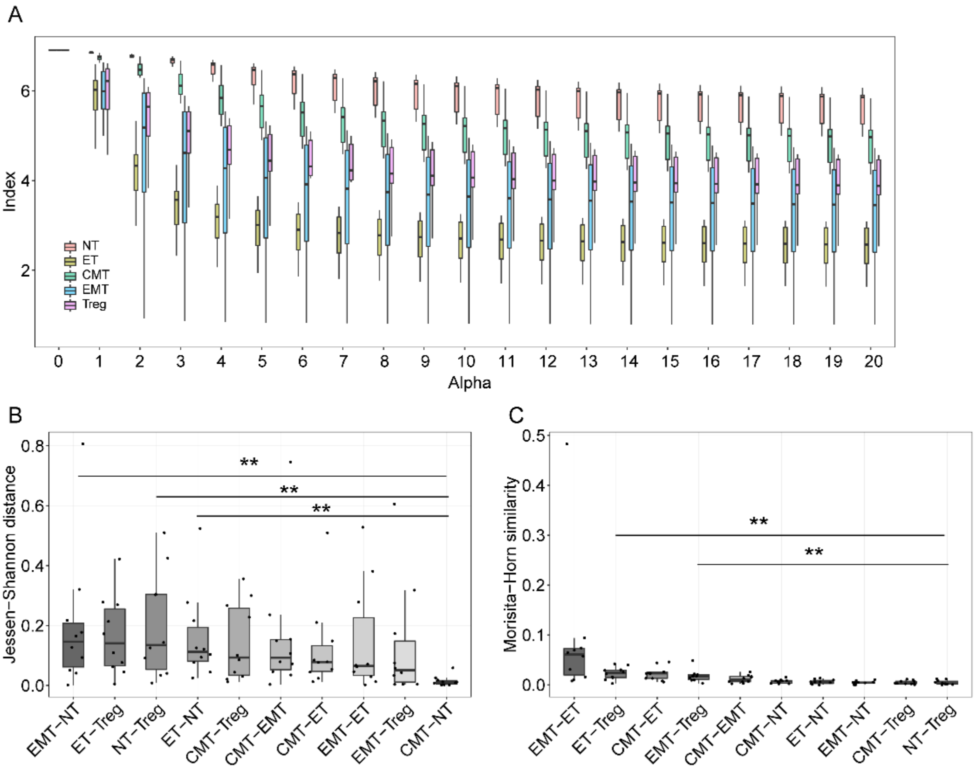


**Supplemental Figure 1. The diversity of TCRB repertoire of five subsets and the relationship of their repertoire architectures.** (**A**) The Renyi entropy index of all subsets with alpha value from 0 to 20. When alpha is equal to 1, the index was calculated as Shannon-index. (**B**) The Jessen-Shannon distance between subsets. The difference between NT and other subsets, and Treg and other subsets were tested. (**C**) The Morisita-Horn similarity between subsets. The difference between NT and other subsets, and Treg and other subsets were tested (Wilcox-ranked test was used, and ** is for p < 0.01).

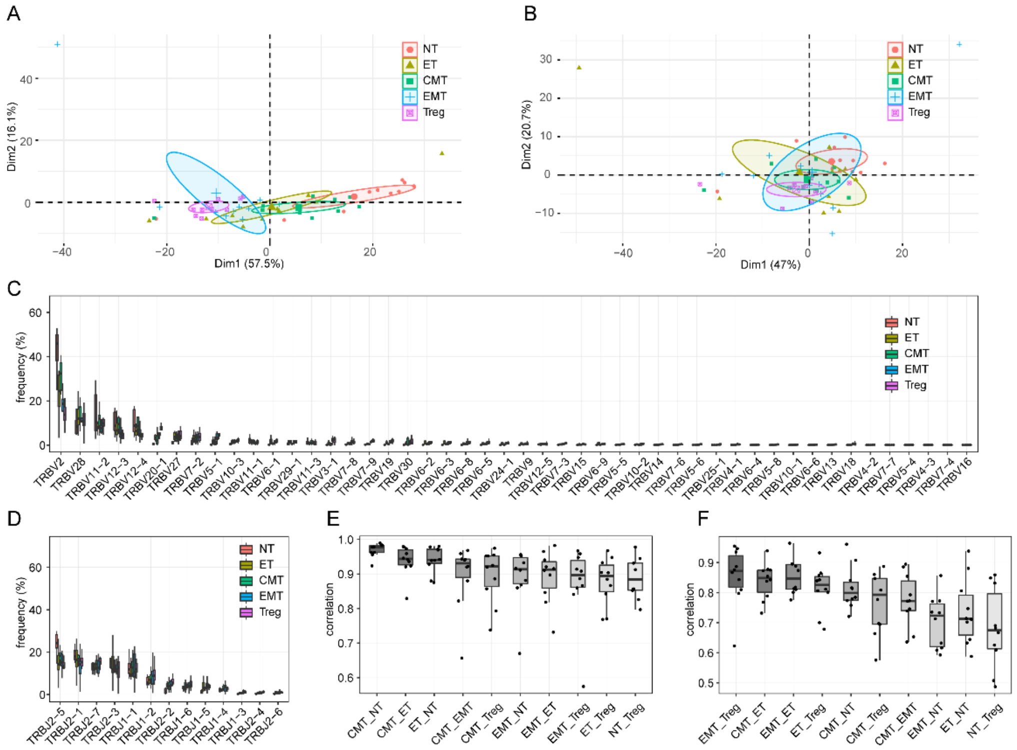

**Supplemental Figure 2. The difference of V- and J-gene usage of subsets and the correlations of gene usage between subsets.** (**A**) PCA based on the frequency of V-genes per subset. (**B**) PCA based on the frequency of J-genes per subset. (**C**) The frequency of V-genes within each subset. (**D**) the frequency of J-gene within each subset. (**E**) The Spearman correlation of V-gene usage between subsets. (**F**) The Spearman correlation of J-gene usage between subsets.

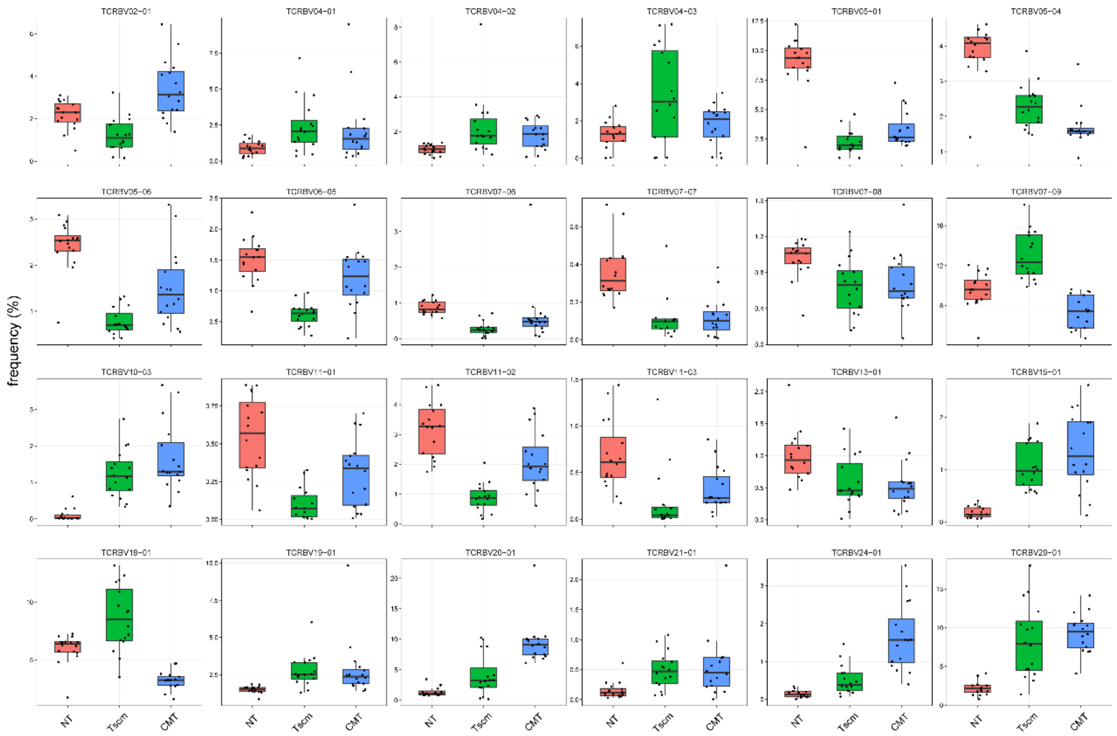


**Supplemental Figure 3. The distinct V-genes used by NT, CMT and Tscm in dataset1.** We examined the V-gene usage between NT and CMT, between NT and Tscm, and then selected distinct genes with a filter of p-value < 0.01 and sample size N > 6. Paired Wilcox-ranked test was used.

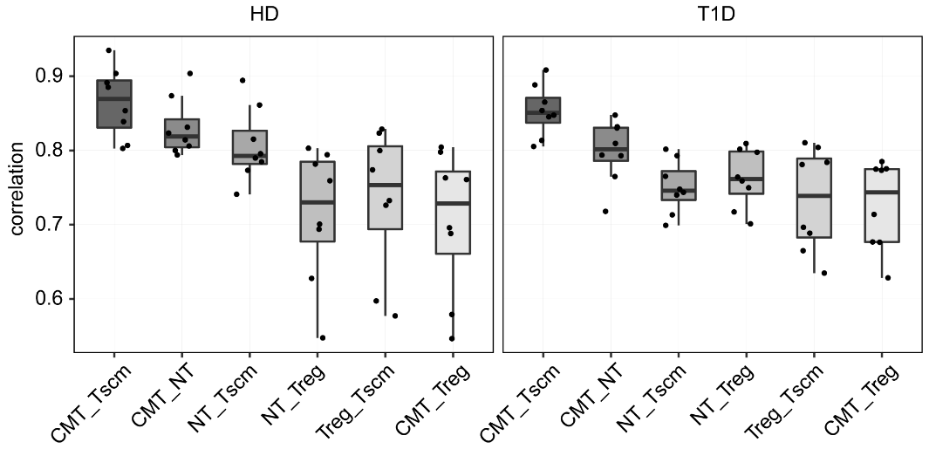


**Supplemental Figure 4. The correlations between subsets were calculated for HD and T1D separately.**

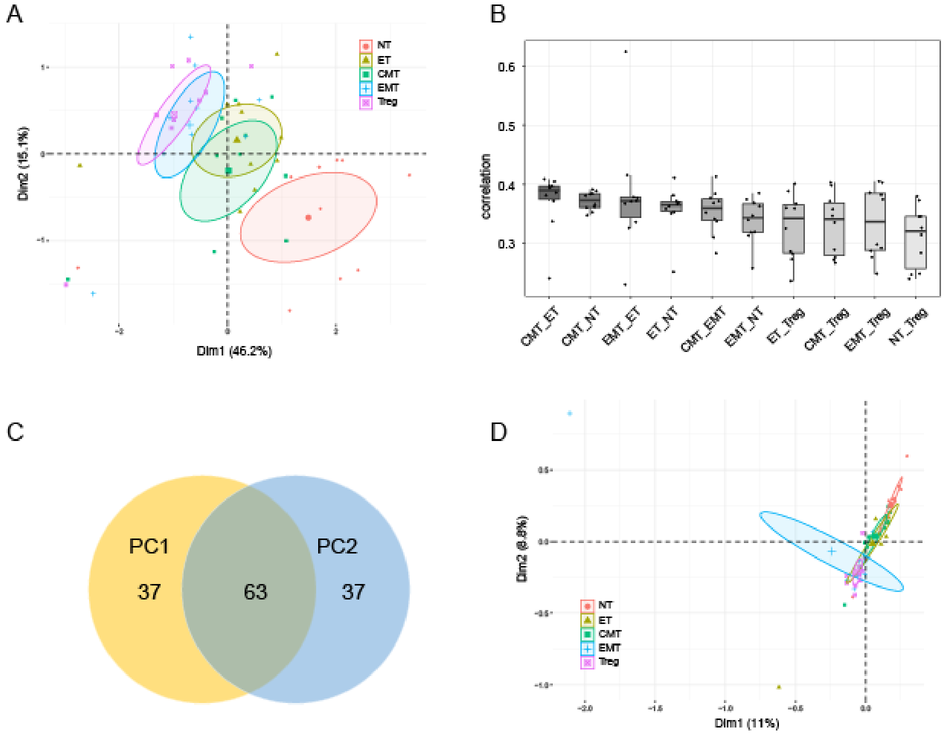


**Supplemental Figure 5. The relationship of sequence composition among subsets in dataset2.** (A) PCA based on the sequence composition for samples. (B) Spearman correlations of k-mer usage between subsets. (C) The overlap of top100 k-mers which mostly contribute to principle component 1 (PC1) and PC2. (D) the PCA based on sequence composition without those located on V- and J-segments for samples.

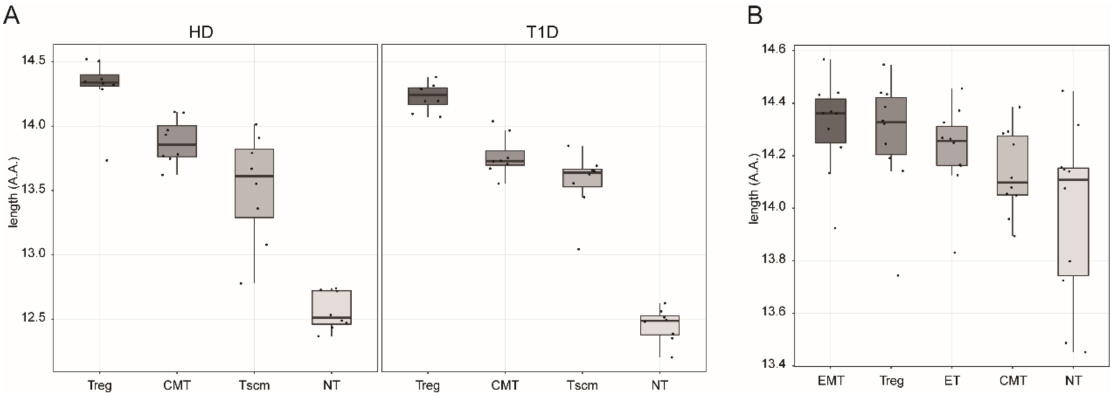


**Supplemental Figure 6. The length distribution of clones within CD4+ subsets.** (**A**) The length distribution of CDR3s of each subset in HD and T1D respectively. (**B**) The length distribution of CDR3s of each subset in RA patients.

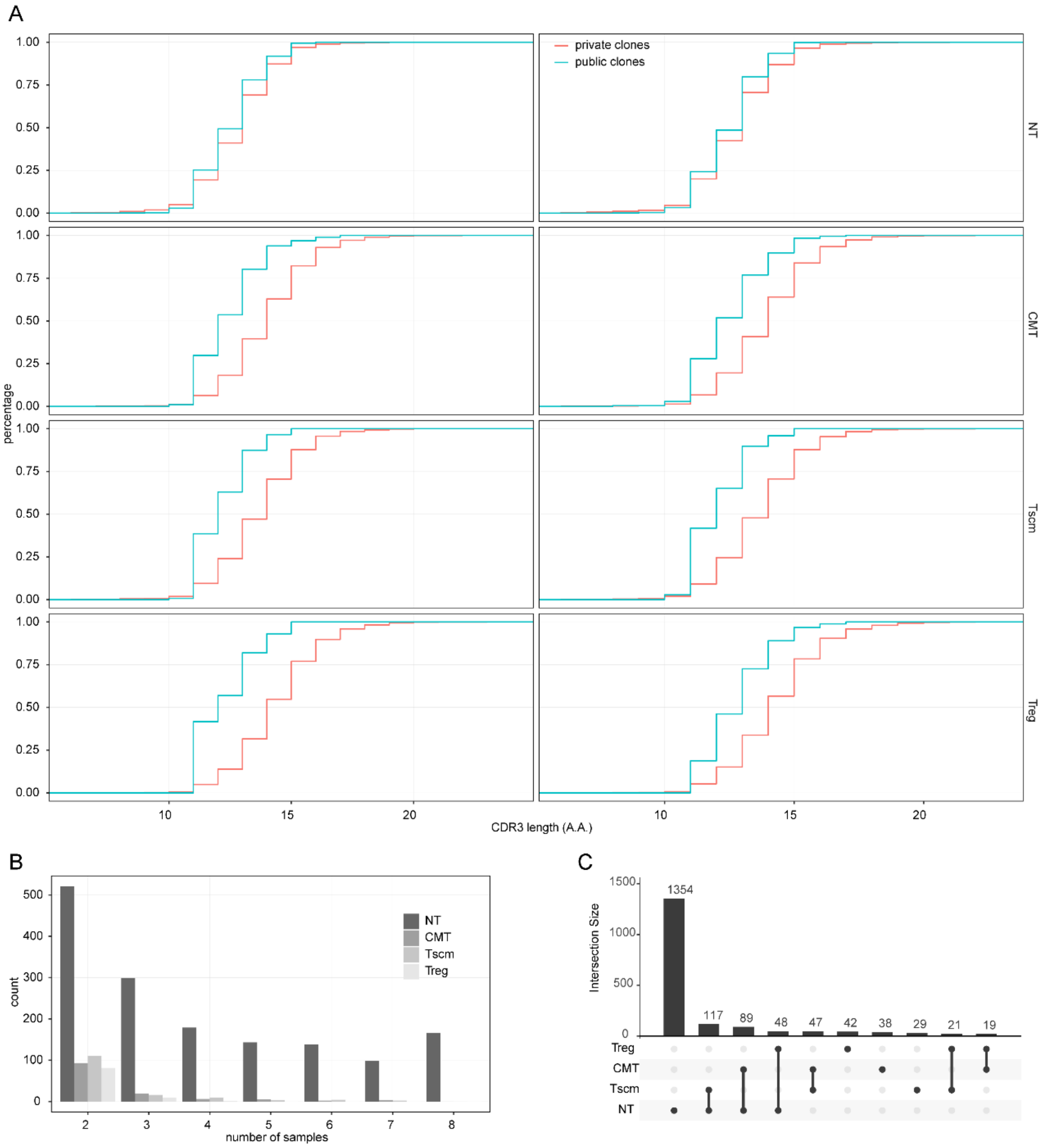


**Supplemental Figure 7. Public clones within each subset.** (**A**) The length distribution of private and public clones within each subset. (**B**)The number of public clones shared by from two to eight T1Ds. (**C**) The overlap of public clones from T1Ds among subsets.

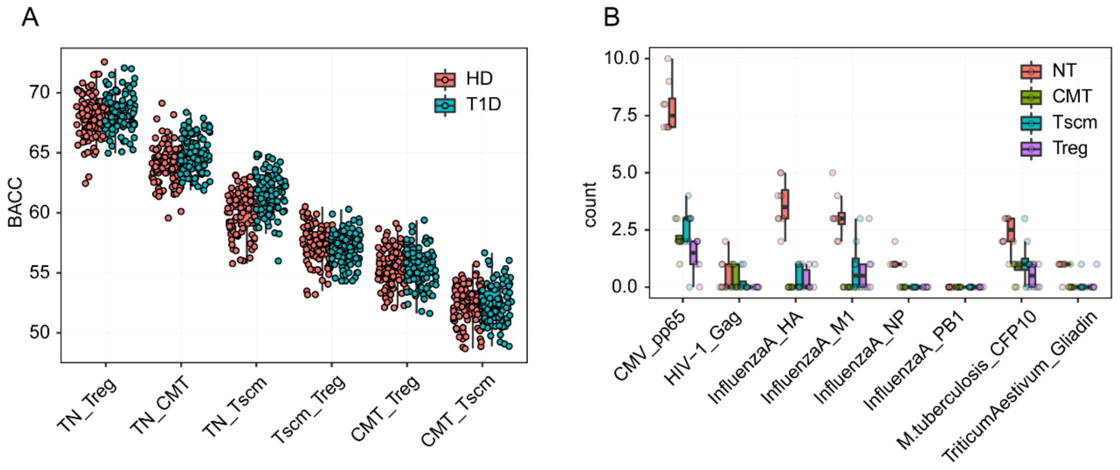


**Supplemental Figure 8. The prediction accuracy of private clones between subsets and the number of clones recognizing given antigen.**
